# Unraveling the Connections between Calreticulin and Myeloproliferative Neoplasms via Calcium Signalling

**DOI:** 10.1101/2021.08.05.455248

**Authors:** Amit Jaiswal, Zhiguo Wang, Xudong Zhu, Zhenyu Ju

## Abstract

Mutations in the form of insertions and deletions (INDEL) in the calreticulin gene lead to essential thrombocythemia which is characterized by the formation of thrombosis. However, the connection between calreticulin INDEL and essential thrombocythemia remains largely elusive. Through combined molecular dynamics simulation and *in-vitro* studies on the wild type and mutated isoforms of calreticulin, the mechanism underlying the calreticulin INDEL induced essential thrombocythemia was investigated at the molecular level. Our results demonstrate that mutations in exon-9 could lead to significant conformational variations of calreticulin structure and thereby reducing its interaction with calcium ions due to decreased electrostatic contributions. The consequence of mutations on calreticulin’s structural integrity was revealed by identifying the key residues and their roles in calcium binding. Furthermore, mutations implemented by CRISPR-Cas9 in exon-9 showed diminished calcium signaling in HEK-293T cells, which agree well with our *in-silico* findings. The current study might help in understanding of the interactions between calreticulin exon-9 INDEL and calcium ions mediated by the structural variations of calreticulin. The results provide useful information for designing novel therapeutic approaches targeting essential thrombocythemia.

## Introduction

Essential thrombocythemia (ET) is a pathological condition which is characterized by the formation of excessive mature blood cells leading to thrombosis [1]. ET belongs to a class of diseases known as myeloproliferative neoplasms (MPNs) which develop through neoplastic transformation of myeloid cells [2]. ET, polycythemia vera (PV), and primary myelofibrosis (PMF) are *BCR-ABL1* negative MPNs, whereas chronic myeloid leukemia (CML) is characterized by the presence of the t(9;22), also known as the Philadelphia chromosome translocation, which results in the generation of the *BCR-ABL1* oncogene [3, 4]. Studies have shown that almost all MPNs are associated with either a *JAK2* V617F mutation or changes in JAK2 signaling, either directly or indirectly [5]. JAK2 drives CML by phosphorylating BCR-ABL1 and triggers 50–60% of ET with myeloproliferative leukaemia virus oncogene (*MPL*) (3–5%) and calreticulin (*CALR*) (30–40%) responsible for the rest of ETs [6, 7]. Though *CALR* mutation has been identified as a marker of MPNs, the underlying molecular process remains largely unrevealed, intriguing us to further explore the mechanism of *CALR*-mediated ETs at the molecular level.

Highly conserved CALR is a calcium-dependent chaperone that is localized in the lumen of endoplasmic reticulum (ER) [8] facilitating calcium (Ca^2+^) homeostasis of the ER by balancing Ca^2+^ communication between ER, plasma membrane, mitochondria and nucleus [9]. Moreover, it has been shown that systemic *CALR* knock-out (KO) is embryonic lethal due to impaired cardiogenesis while a conditional KO perturbs secondary messengers such as inositol 1,4,5-trisphosphate [10]. Structurally, CALR contains three domains: a global domain called N-domain responsible for chaperone-like functions, a proline-rich P-domain containing a flexible arm-like structure, and a C-terminal acidic domain comprising of a KDEL motif responsible for an ER retention signal [11, 12]. The KDEL sequence acts as a signature in other ER-resident chaperons to recognize and retrieve KDEL-containing proteins from ER-Golgi transport and *vice versa* [13].

Calcium homeostasis is maintained in the cell involving various calcium transport molecules, buffers and sensors that control free and bound calcium levels in different cellular compartments such as cytoplasm, nucleus, mitochondria and ER [14]. Apart from protein folding and transportation, the ER primarily functions as a calcium storage pool as many calcium related proteins reside on the ER membrane [15]. Accumulation of misfolded proteins in the ER can alter the calcium balance resulting in the development of many diseases [16, 17]. Changes in the calcium concentration are regulated by extracellular calcium uptake or intracellular calcium release in order to initiate a calcium electrochemical gradient flow resulting in calcium signaling [18]. There are two groups of calcium-binding proteins, namely cytosolic and organellar which show a different pattern of affinity towards calcium binding. The cytosolic group binds calcium with high affinity and low capacity while the organellar group binds it with high capacity and low affinity [19]. All these processes are highly orchestrated in a very intricate manner and if disturbed, serious consequences may occur such as cardiac arrhythmia [20].

Although biochemical studies have provided plenty of information about *CALR* [5, 21, 22], very little is known about the C-domain, especially the molecular mechanism of the C-terminus related to calcium ions due to the lack of structural information. Here we aim to elucidate the molecular mechanism of *CALR* in the context of calcium homeostasis and ET by adopting structural bioinformatics and CRISPR-Cas9 approaches. Our molecular dynamics (MD) study suggests that loss of *CALR* C-terminal residues would disrupt the negative potential of exon-9 leading to inefficient calcium binding. We propose that calcium-overload is supposed to trigger an aberrant calcium signaling pathway, leading to persistent activation of platelets culminating in the clinical manifestation of thrombosis. Exon-9-mutant HEK-293T cells exhibited diminished cellular calcium fluorescence compared to WT cells, strongly supporting our computational findings. Overall, the current work provides mechanistic insights in the binding of *CALR* and calcium ions and also novel information for developing new therapeutics targeting ET.

## Results

### Sequence and Structural Characterization of Calreticulin

The gene locus of *CALR* is situated on chromosome 19 and is 417 amino acids long consisting of nine exons (Figure 1A and B). Functional domain analysis by Pfam revealed the presence of a single family present in two regions of calreticulin protein: one starting from 1 to 253 and the other one from 293 to 367 while Interproscan disclosed two protein families, namely the Glucanase domain family comprising residues 19 to 217 and a calreticulin protein family beginning from amino acid 201 to 315. The Uniprot web-server showed the presence of three domains: a N-domain comprising amino acids 18-197, a P-domain from 198-308 and a C-terminal acidic domain from 309 to 417 as shown in Table 1 and Figure 1C.

**Figure1:**
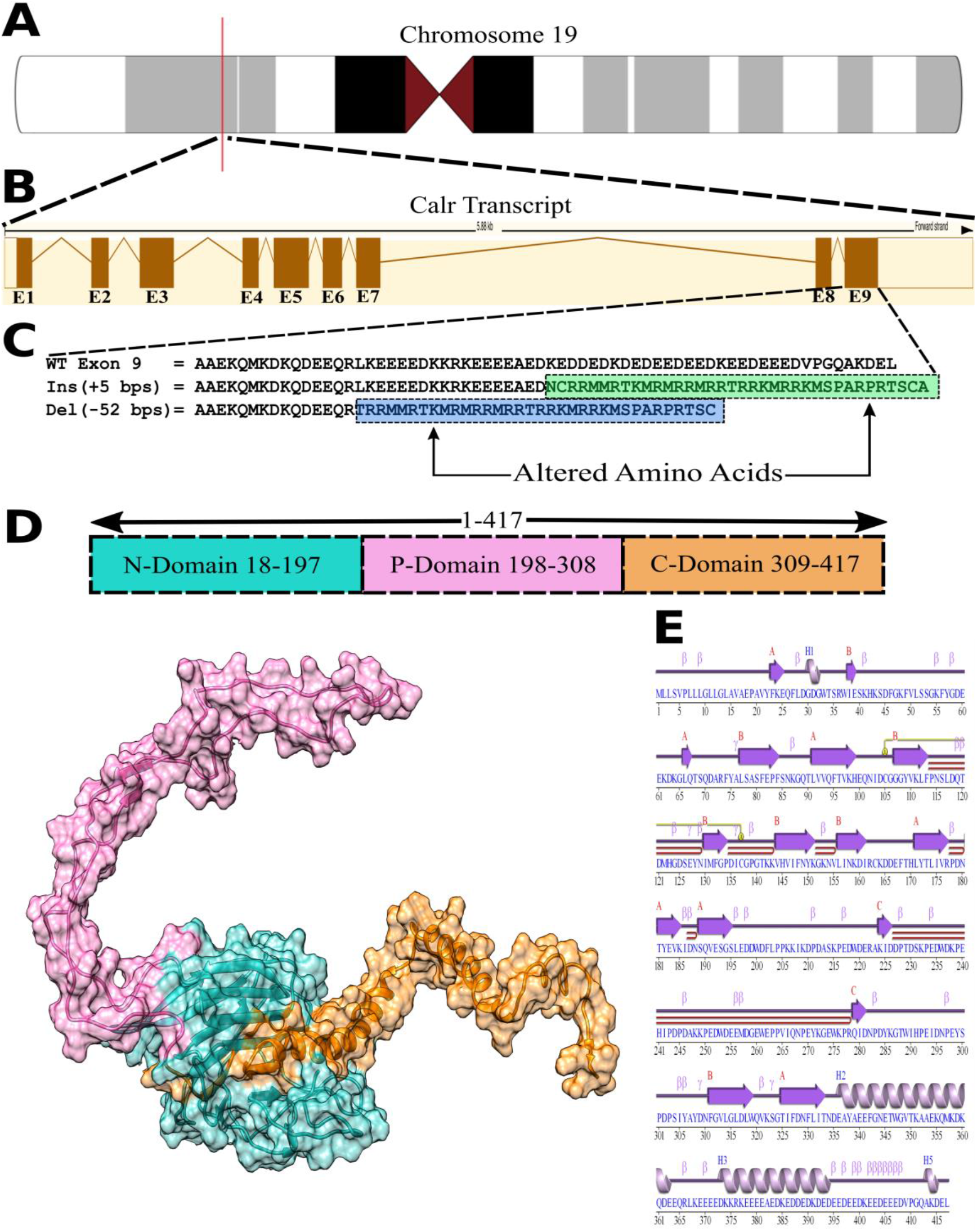
Gene locus transcript, sequence and structural information of calreticulin (CALR). (A) Gene locus information of *CALR*. (B) Transcript information showing presence of nine exons. (C) differences in the amino acid sequences between WT and mutants. Ins gains 5 bps in Exon-9 while Del loses 52 bps from exon-9. Altered amino acid sequence in Ins is highlighted in green color while in blue color in Del. (D) 3D structure of *CALR*. The domains are coloured in different colours. N or globular domain is colored in sky-blue, P-domain in pink, C-domain in orange. (E) Secondary structure of *CALR*. PDBsum webserver was used to prepare the figure.

**Table 1.**
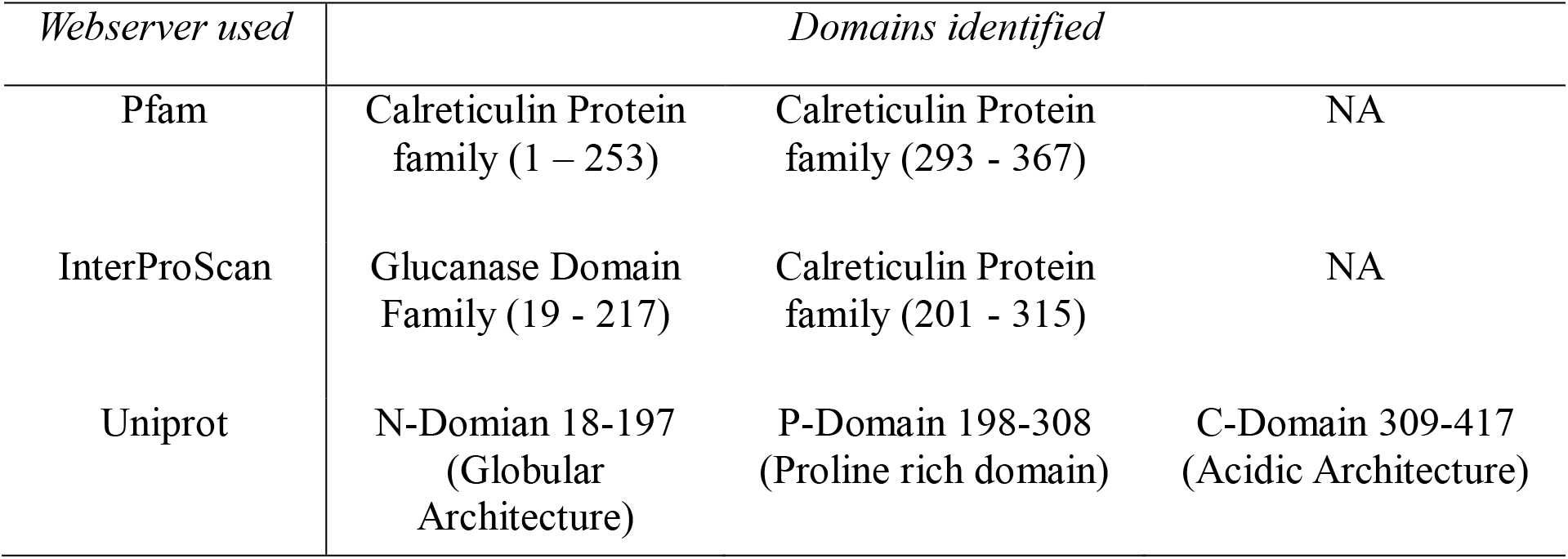
List of protein domains identified by Pfam, InterProScan and Uniprot web server

| *Webserver used* | *Domains identified* |  |  |
| --- | --- | --- | --- |
| Pfam | Calreticulin Protein family (1 – 253) | Calreticulin Protein family (293 - 367) | NA |
| InterProScan | Glucanase Domain Family (19 - 217) | Calreticulin Protein family (201 - 315) | NA |
| Uniprot | N-Domian 18-197<br>(Globular Architecture) | P-Domain 198-308<br>(Proline rich domain) | C-Domain 309-417<br>(Acidic Architecture) |

Multiple Sequence alignments of *CALR* to its homologous sequences in various other mammalian species using the Clustl-X algorithm showed the existence of highly conserved amino acids throughout the sequence with an exception of the C-terminal region in exon-9 leading to a charge variation. The last seven amino acids comprising of the KDEL motif in exon-9 were also highly conserved portraying the importance of this region (Figure. S1). Lack of structure for full-length sequence in the protein data bank required us to use the I-TASSER web-server to generate full-length structures, reasoning that full-length models would provide a more complete understanding of all domain movements and dynamics arising due to mutations. In order to validate the structures, we performed a Ramachandran plot assessment which suggested that 80.3% of residues were present in the most favored region for the WT while 81% and 82% for Ins and Del variants/mutations, respectively (Figure S2). The structural quality score provided by Verify-3D proposed 85.37%, 80%, 83.75%, and ERRAT proposed 69.14%, 82.59 %, 76.70% for WT, Ins and Del, suggesting the reliability of the I-TASSER models for further studies (Table2).

**Table 2.** Structural Validation of generated 3D structures through Ramachandran plot assessment, Verify 3D and ERRAT scores

|  | 3D Structural Assessment |  |  |
| --- | --- | --- | --- |
|  | WT | Ins | Del |
| Residues in favored region (%) | 80.3 | 81 | 82 |
| Residues in additionally allowed region (%) | 17.7 | 16.5 | 16.6 |
| Residues in generously allowed region (%) | 1.7 | 1.4 | 1.2 |
| Residues in disallowed region (%) | 0.3 | 1.1 | 0.3 |
| Verify _3D score (%) | 85.37 | 80 | 83.75 |
| ERRAT score (%) | 69.14 | 82.59 | 76.70 |

The selected models were then subjected to molecular dynamics simulation (MDS) using the GROMACS 4.5.6 package. All proteins: WT, Ins and Del were simulated in the presence and absence of calcium ions in order to decipher the structural changes governed by the effects of calcium. The parameters for the simulation system are stated in Table S2. A preliminary study of secondary structure using the PDBsum web-server gave a discernment about secondary structures. The number of strands, helices, helix-helix interactions and turns was higher in Del and Ins while β-turns were higher in the WT (Table S3). However, when WT-Ca was compared to Ins-Ca and Del-Ca, a higher number of β-strands and β-turns was found while a lower number of helices was observed in WT-Ca as compared to other forms (Table S3). The minimum and maximum potential energy (PE) was observed to be −2732658 kJ/mol and −2720337 kJ/mol for WT, −3103612 kJ/mol and −3087532 kJ/mol for Ins and −3664817 kJ/mol and −3650847 kJ/mol for Del counterparts while the kinetic energy (KE) analysis showed contrasting results: The WT had the lowest KE compared to mutated equivalents (Table S4). Pressure and enthalpy were higher in the WT compared to mutated counterparts as can be seen from Table S4.

### Mutations in Exon-9 display huge structural Aberrations in the C-Domain

Three-dimensional structures of WT, Ins and Del were significantly dissimilar despite of the fact that all the proteins under investigation are identical from exon-1 to exon-8. Mutated isoforms such as Ins and Del had a similar surface structure where the C-terminal region collapsed to form a continuous loop-like structure. The two helices in exon-9 overlap with each other while it is stretched out in an elongated conformation in the WT (Figure 2 A-C). To measure the conformational stability of the proteins, Root Mean Square Deviation (RMSD) profiles for protein backbone residues were generated. RMSD results presented in figure 2D show the trajectory of the WT to be stable after 42ns whereas for Ins it took around 40ns while Del was unable to attain proper stability even after 50ns. The radius of gyration (Rg) was calculated by comparing the protein backbone residues with the native structure in order to observe the compactness of the protein structures. The average Rg for the WT was 3.45nm while that of Ins and Del was 2.79nm and 2.54nm, respectively (Figure 2E). Residue fluctuation for each protein was also calculated throughout a 50ns timescale by computing Root Mean Square fluctuation (RMSF) as can be seen in figure 3A. RMSF results demonstrated the WT to have a huge fluctuating P-domain and a C-terminal domain ranging from 230-280 (0.5-2.8 nm) to 370-417 (1-2.3 nm) while in Ins the residue range was reduced between residues 210-290 (0.4-1.2 nm) and 370-420 (0.6-1.1 nm). In Del residues 220-290 showed an overall fluctuation which ranged from 0.6-1.2 nm. Together, given the results from RMSD, Rg and RMSF; the WT proved to be highly unstable, most likely because of the presence of an unstable P-domain and C-domain.

**Figure 2:**
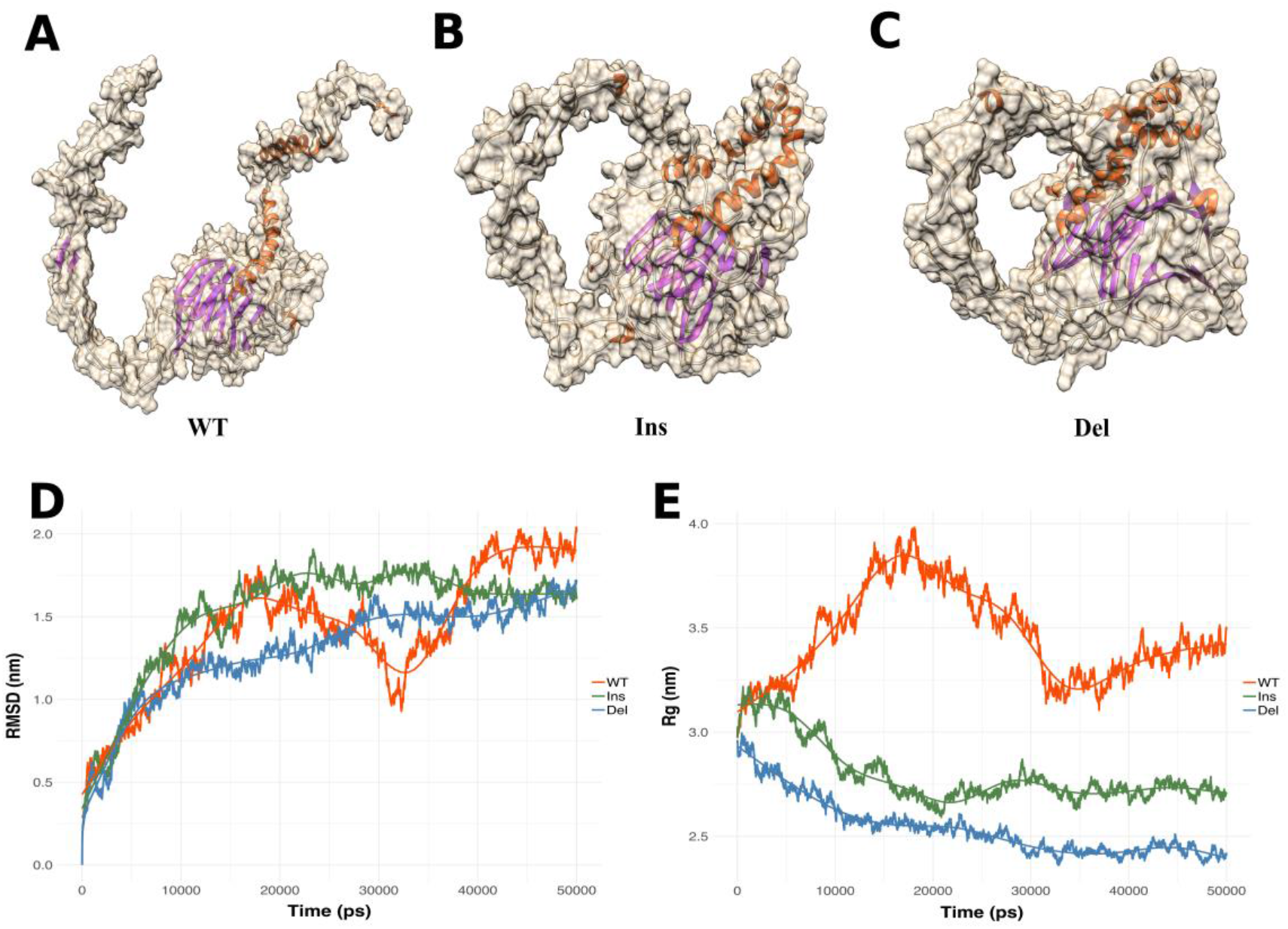
Comparison of WT, Ins and Del based on structure, RMSD and Rg in absence of calcium. (A), (B) and (C) represent 3D structures of WT, Ins and Del, respectively. The cartoon inside the surface view represents the secondary structure. Alpha-helices are colored in orange, beta-sheets in magenta. (D) Root mean square deviation (RMSD) of WT, Ins and Del in red, green and blue, respectively. (E) Radius of Gyration (Rg) of WT, Ins and Del (red, green and blue, respectively).

**Figure 3:**
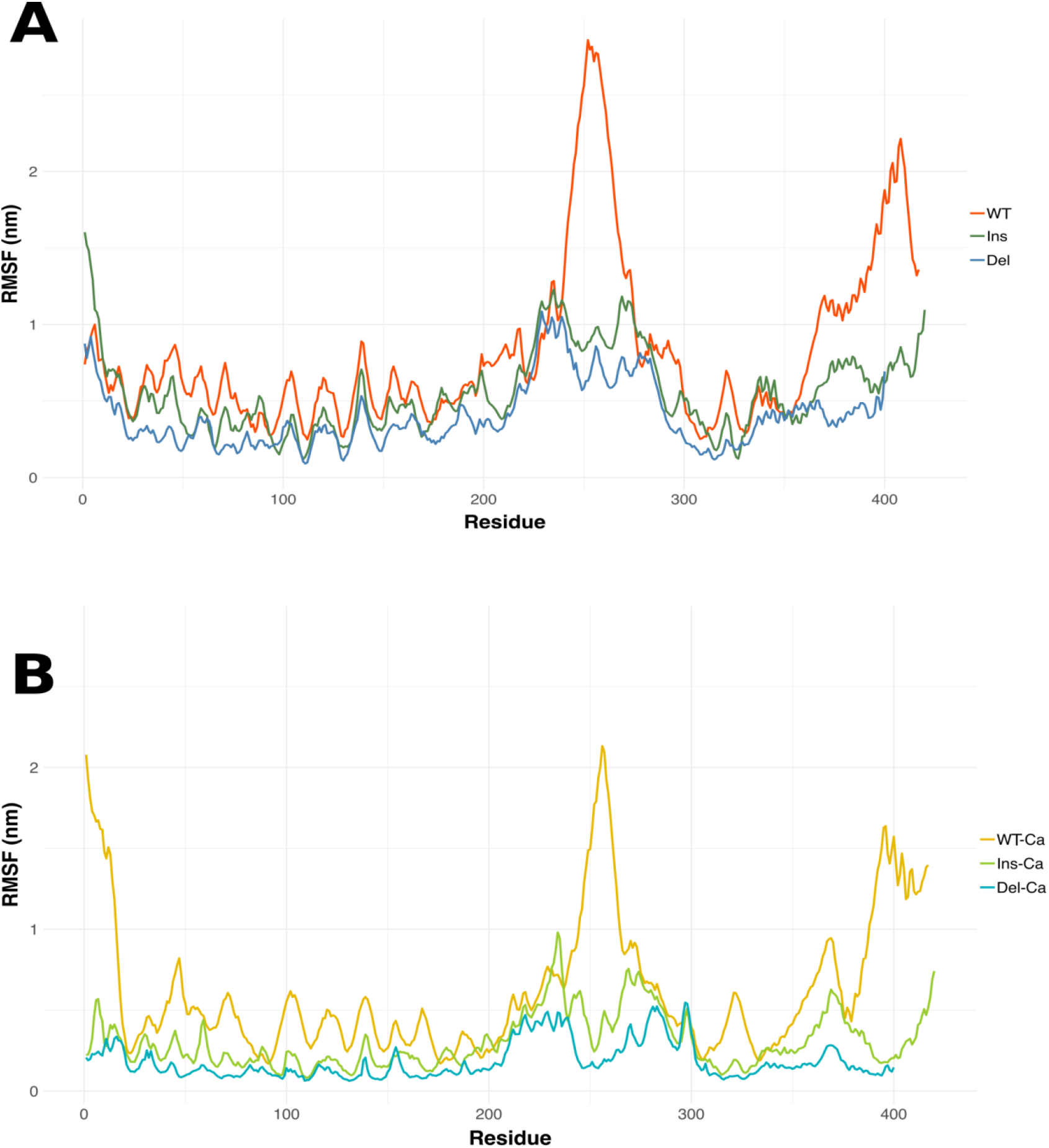
Comparison of RMSF in presence and absence of calcium ions where higher peak indicates higher residue fluctuations. (A) Root Mean Square Fluctuations (RMSF) in the absence of calcium. Red, green and blue color represent WT, Ins and Del, respectively. (B) RMSF in the presence of calcium. Golden, light-green and sky-blue represent WT-Ca, Ins-Ca and Del-Ca, respectively.

Structures of WT and mutated genes (Ins and Del) after 50ns of simulation were structurally compared using structural superimposition which showed huge structural differences (Figure 5). The C-terminal region comprising of the KDEL motif of WT had an open form compared to mutated equivalents. The hinge region comprising of R366 – E369 in the C-domain of the WT had an open α-helix conformation while in the mutated forms it was closed. The presence of a β-sheet was also visible in the extended arms of the P-domain ranging in the region K224-D226 and Q279-D281, respectively. Three-dimensional structural analysis showed that the N-domain or the globular domain fluctuated less compared to the P- and C-terminal acidic domains in all the proteins (Figure 5A,B). A Root Mean Square Deviation (RMSD) score of 21.776 Å was observed between WT and Ins while 25.505 Å between WT and Del (Figure4).

**Figure 4:**
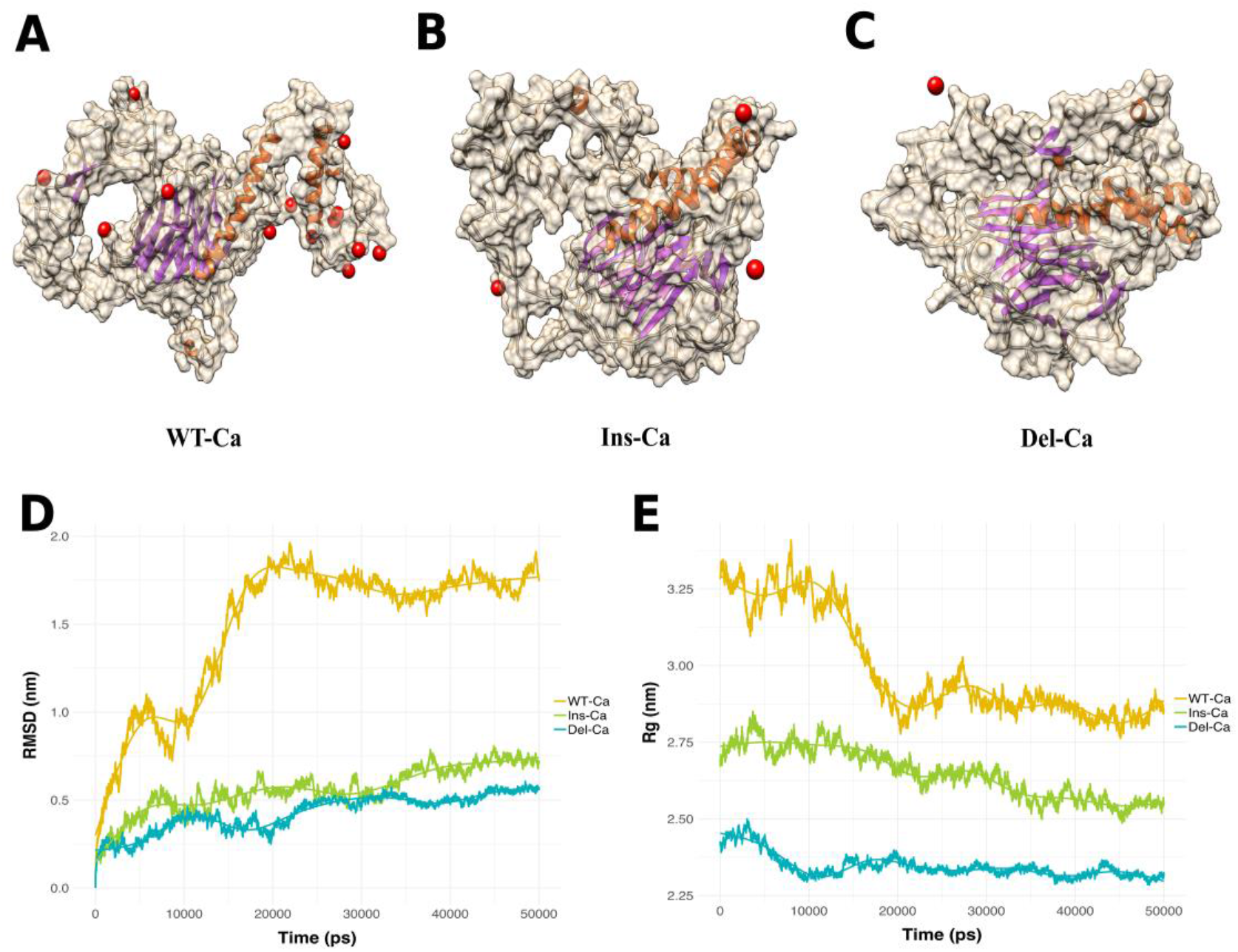
Comparison of WT-Ca, Ins-Ca and Del-Ca based on structure, RMSD and Rg in presence of calcium. (A), (B) and (C) represent 3-D structures of WT-Ca, Ins-Ca and Del-Ca. The cartoon representation inside the surface view is based on secondary structure. α-Helices are colored in orange, β-sheets in magenta. (D) RMSD of WT-Ca, Ins-Ca and Del-Ca colored in golden, light-green and sky-blue. (E) Rg of WT-Ca, Ins-Ca and Del-Ca colored in golden, light-green and sky-blue.

**Figure 5:**
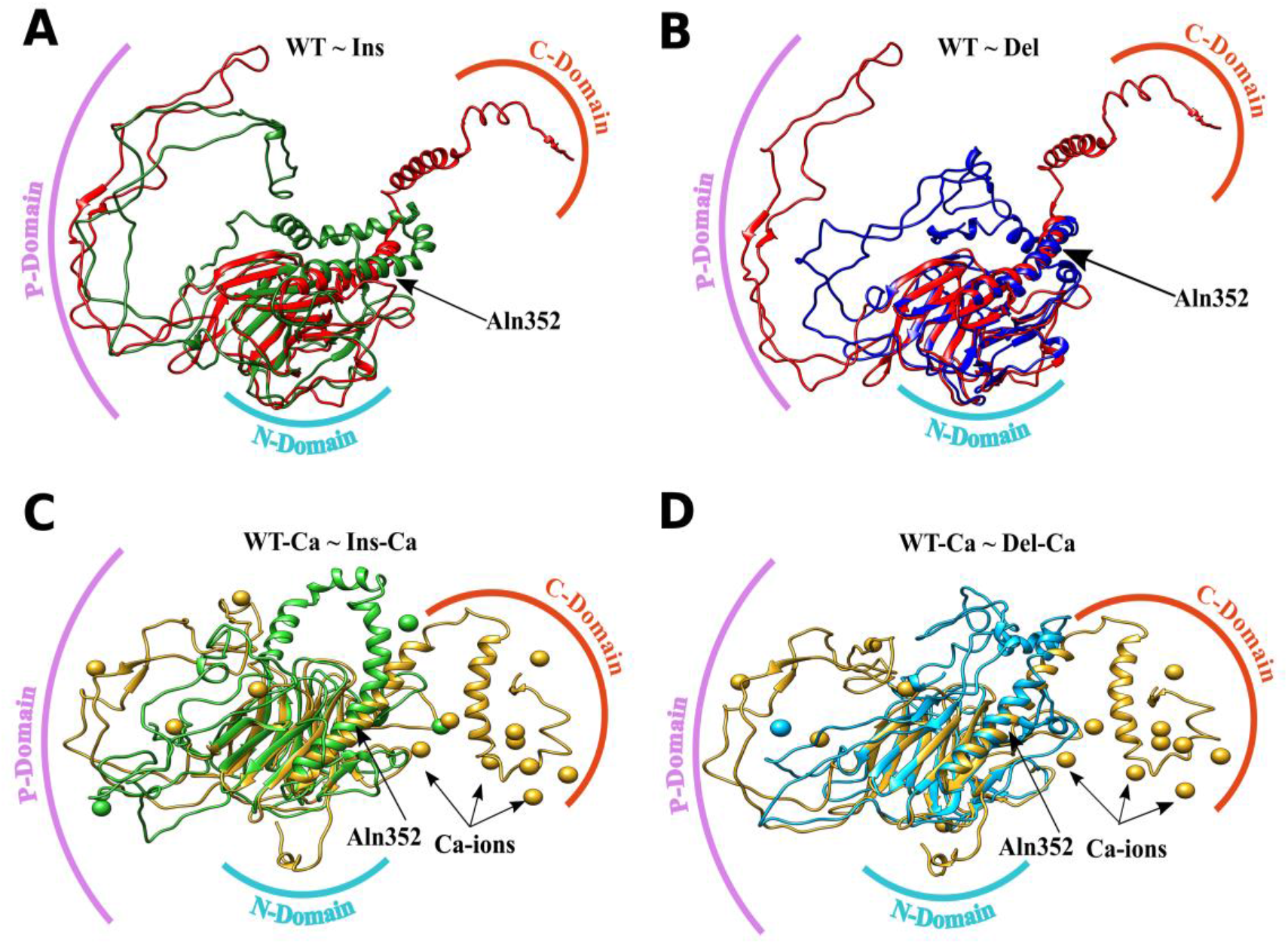
Structural alignment between proteins in presence and absence of calcium ions. (A) and (B) structural alignment of WT with Ins and Del in absence of calcium ions. (C) and (D) the same as in (A) and (B) but in presence of calcium ions. WT (in red), Ins and Del are colored in green and blue. WT-Ca, Ins-Ca and Del-Ca are colored in golden, light-green and sky-blue. The starting residue for exon-9 is indicated as Aln352. Various protein domains are colored in various colors. The P-domain is indicated in pink, N-domain in blue and C-domain in orange color while calcium ions are represented as golden balls.

We then aimed to analyse the secondary structure formation through the simulation of 50ns and thus we performed DSSP (Define Secondary Structure of Proteins) by using the inbuilt DSSP tool in the GROMACS package. The DSSP results for WT, Ins and Del showed a significant difference. Residues 413 to 416 attained 3-helix configurations throughout the simulation while Ins and Del lost this configuration. Moreover, in WT residues 397 to 409 took a configuration of a β-turn after 40ns of a MD run and remained stable throughout the simulation, while in Ins and Del the aforementioned residues took a configuration of α-helices and coils, respectively (FigureS3). The majority of residues ranging between 350-395 in the WT were an α-helix and the same holds true for Ins and Del. However, Del formed an additional 5-helix structure from 380-384 indicating a structural difference from WT. The most crucial point of observation was that exon-9 residues which ranged between 396-410 and 413-416 showed the formation of a β-turn and a 3-helix throughout the simulation which was not present in both mutated proteins. We also observed differences in the residues ranging from 311-335 having β-sheet conformation in the WT but this had been greatly reduced in mutated proteins (Figure S3).

### The Presence of Ca^++^ ions provides structural stability

In the presence of calcium, the WT-Ca showed lower PE and higher KE compared to Ins-Ca and Del-Ca. (Table S4) while structural analysis revealed that the P- and C-domains held their unique structure quite firmly surrounded by calcium ions in the WT. In case of Ins-Ca and Del-Ca, the P- and C-domains merged together losing their structural symmetry. In addition, the number of calcium ions that interacted with the C-terminus was much less in mutated proteins compared to the WT (Figure 4 A-C). RMSD analysis of protein backbone revealed that WT-Ca took only about 20ns to become stable and remained stable all the way through. In contrast, Ins-Ca and Del-Ca attained stability at around 45ns evidencing that WT in the presence of calcium had more firmness and stability compared to its mutated equivalents (Figure 4D).

Furthermore, WT-Ca had less structural integrity in comparison to Ins-Ca and Del-Ca, possibly because of non-functional disintegrated N- and C-terminal domains which remain intact in the mutants. The average Rg for WT-Ca was 2.99 nm while that of Ins-Ca and Del-Ca was 2.65 nm and 2.3 nm (Figure4E). RMSF results for WT-Ca ranged from residues 220-280 (0.5-2.1nm) to 370-417 (0.3-1.6 nm) while for Ins-Ca and Del-Ca the fluctuating residues resided between 220-290 (0.5-0.9 nm) and 210-300 (0.3-0.5 nm), respectively (Figure 3B). The same was found in the region of 370-417 except for the fact that in WT-Ca the fluctuations declined severely. Taken together with the results from RMSD, Rg and RMSF, we concluded that the majority of calcium ions anchors to the C-terminal region and imparts stability.

Moreover, 3D structural imposition showed a deviation of 25.172 Å between WT-Ca and Ins-Ca, while it was 20.776 Å between WT-Ca and Del-Ca. These huge deviations suggest that calcium ions play a significant role in stabilizing the P- and C-domains significantly in WT (Figure.5 C-D). For the DSSP results, the WT showed similar signatures to its non-calcium counterparts except for an additional formation of a 3-helix in the region of 399-402, while in Ins-Ca and Del-Ca the majority of configurations was similar to the non-calcium counterparts. In addition, the residues ranging from amino acids 154-167 in Ins-Ca showed a coil configuration while WT-Ca revealed a β-sheet. Moreover, residues 153-155 showed a β-turn while residues 160-162 only formed β-sheets and the remaining residues all formed exclusively coils (Figure S4).

### Ca^++^ ions but not Na^+^ show more binding Affinity towards the C-terminal Region

To determine structural differences, we compared the WT structures both in the presence and absence of calcium. The WT in a calcium environment attained stability just after 20ns while the WT in the presence of Na^+^ took 45ns. Furthermore, the average Rg of WT-Ca was about 2.9 nm which remained stable while that of WT ranged between 3-4 nm and had yet to attain stability (Figures 2, 3 and 4). Finally, the RSMF results suggested the P-domain to be less fluctuating in WT-Ca than WT. Moreover, the C-terminal region comprising of the KDEL motif also showed a huge decline in fluctuation indicating the region to be more stable in the presence of calcium (Figure. 3). Furthermore, structural superposition of protein structures showed huge variations between WT and WT-Ca with a deviation of 23.41Å especially in the P- and C-domains (Figure 6). Comparing the DSSP results for calcium equivalents, the WT showed similar signatures to WT-Ca with an exception by forming a 3-helix in the region of 399-402 (Figure S3). The secondary structures of Ins-Ca and Del-Ca were similar to the non-calcium counterparts suggesting that the presence of calcium could not disturb the structural stability in mutated proteins (Figure. S3 and S4). Residues 208-209 and 293-294 in WT-Ca underwent conformational changes forming a β-bridge to a β-sheet which was absent in mutated proteins.

**Figure 6:**
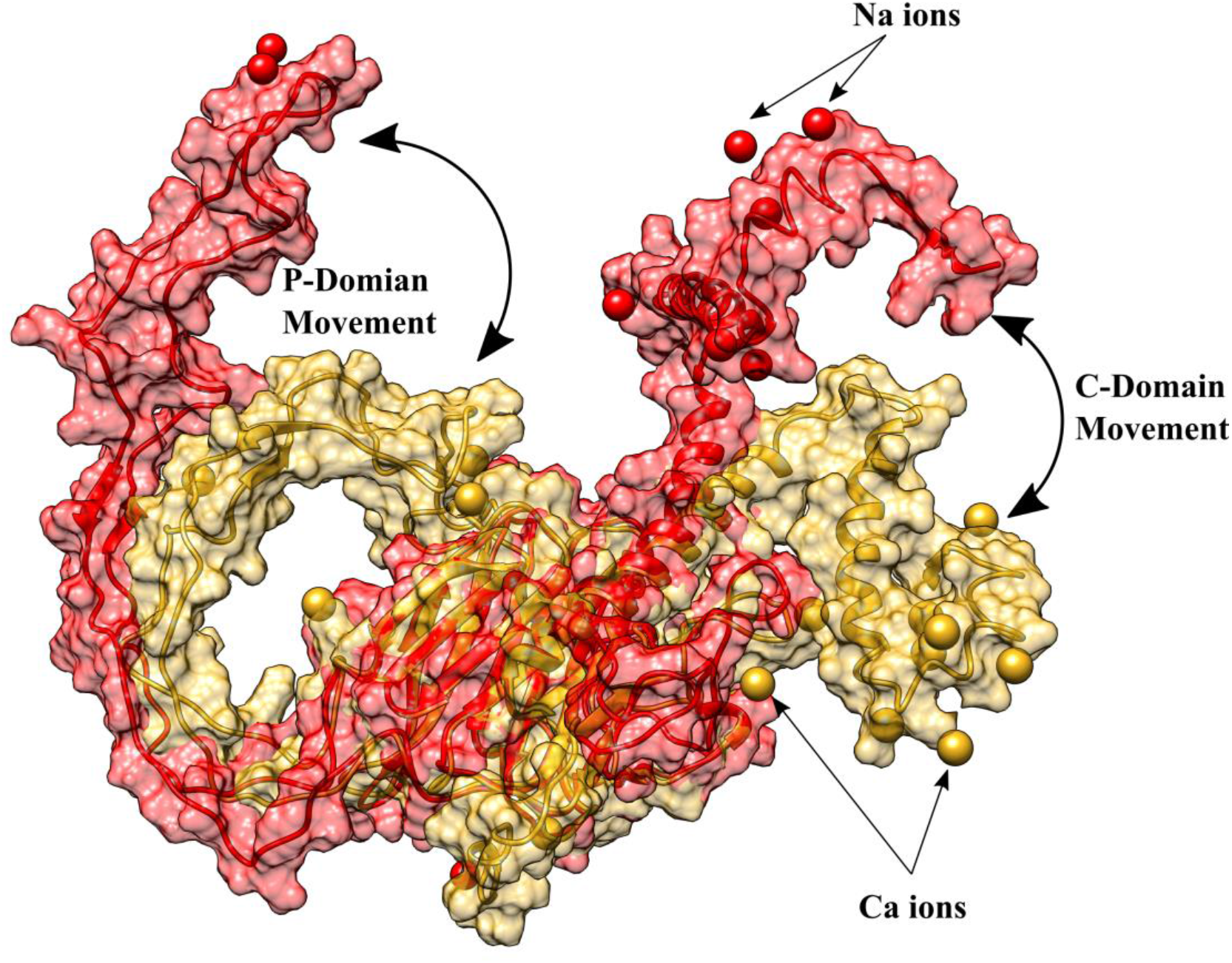
Structural changes of WT in presence and absence of calcium ions. 3D structure of WT in absence of Ca^++^ ions is shown in Red while 3D structure of WT in presence of Ca^++^ ions (golden.) Both structures were superimposed to each other. The structural variation and movement of P and C-domain are represented by arrows.

The total number of amino acids present in exon-9 in the WT is 66 while in Del and Ins it is 49 and 69, respectively. The Protparam tool indicated that 75%, 38% and 18% of amino acids were negatively charged in the WT, Ins and Del counterparts suggesting that mutation (INDEL’s) lead to the loss of negative amino acids thereby decreasing the entire negative potential (Figure S5A). The C-terminal region was highly unstable as evidenced by the Instability Index and IUPred2 webserver [44] (Figures S5B and S5C). The percentage of distinct amino acids in exon-9 also varied a lot. The amount of amino acids like aspartic acid, glutamine, leucine, glycine and valine was significantly higher in the WT compared to mutants while levels of arginine, methionine and cysteine and proline were lower in the WT than in its mutated counterparts (Figure S5D) and the isoelectric point of exon-9 was much lower in the WT compared to the other two forms (Figure S5E). However, analysing the aliphatic index, we found that the WT protein had a much higher presence of aliphatic side chains (alanine, valine, isoleucine, and leucine), signifying an increase in the thermo-stability of exon-9 [45]. Atomic composition of exon-9 in the WT showed the presence of sulphur to be around 5 % while in the mutational counterparts it was 45% each proposing the mutated proteins to be more susceptible to forming thiol (S-H) and disulfide bonds (S-S) (Figure S5F). It has been reported that methionine and cysteine are the only two natural amino acids which have sulphur atoms [46]. Thus, it appeared that the mutated proteins had a higher number of methionine and cysteine amino acids in exon-9 compared to the WT.

We then explored the calcium dynamics of calreticulin in the presence of exon-9. The Mindist tool from the Gromacs package informed us about the distance of calcium ions from exon-9 throughout the simulation. The average minimum distance of calcium ions from exon-9 was found to be 3.7 Å in WT-Ca and remained stable, while for the WT it was very chaotic as the variations ranged from 2 - 4 Å and were unstable throughout the simulation (Figure7A). We also performed a H-bond analysis using the Gromacs tool to determine the difference between intra-molecular H-bonds within exon-9. To our surprise, a higher number of H-bonds was observed in the presence of calcium ions suggesting that calcium ions provide structural stability to the exon-9 region of *CALR* (Figure 7B). Moreover, it is worth to note that Na^+^ ions were unable to provide enough stability to balance the negative amino acids present in exon-9 indicating the importance of calcium ions. In order to study the electrostatic differences between the WT and mutated proteins, Adaptive Poisson-Boltzmann Solver (ABPS) studies were conducted using Pymol software. The presence of calcium supported the folding of the C-terminal acidic domain and this fold remained stable throughout the MD run through increased electrostatic interaction. The electrostatic charges in mutated proteins were less due to mutations in exon-9 while the presence of calcium influenced both N-and C-termini by increasing their electrostatic potential (Figures 7 C, D and S6).

**Figure 7:**
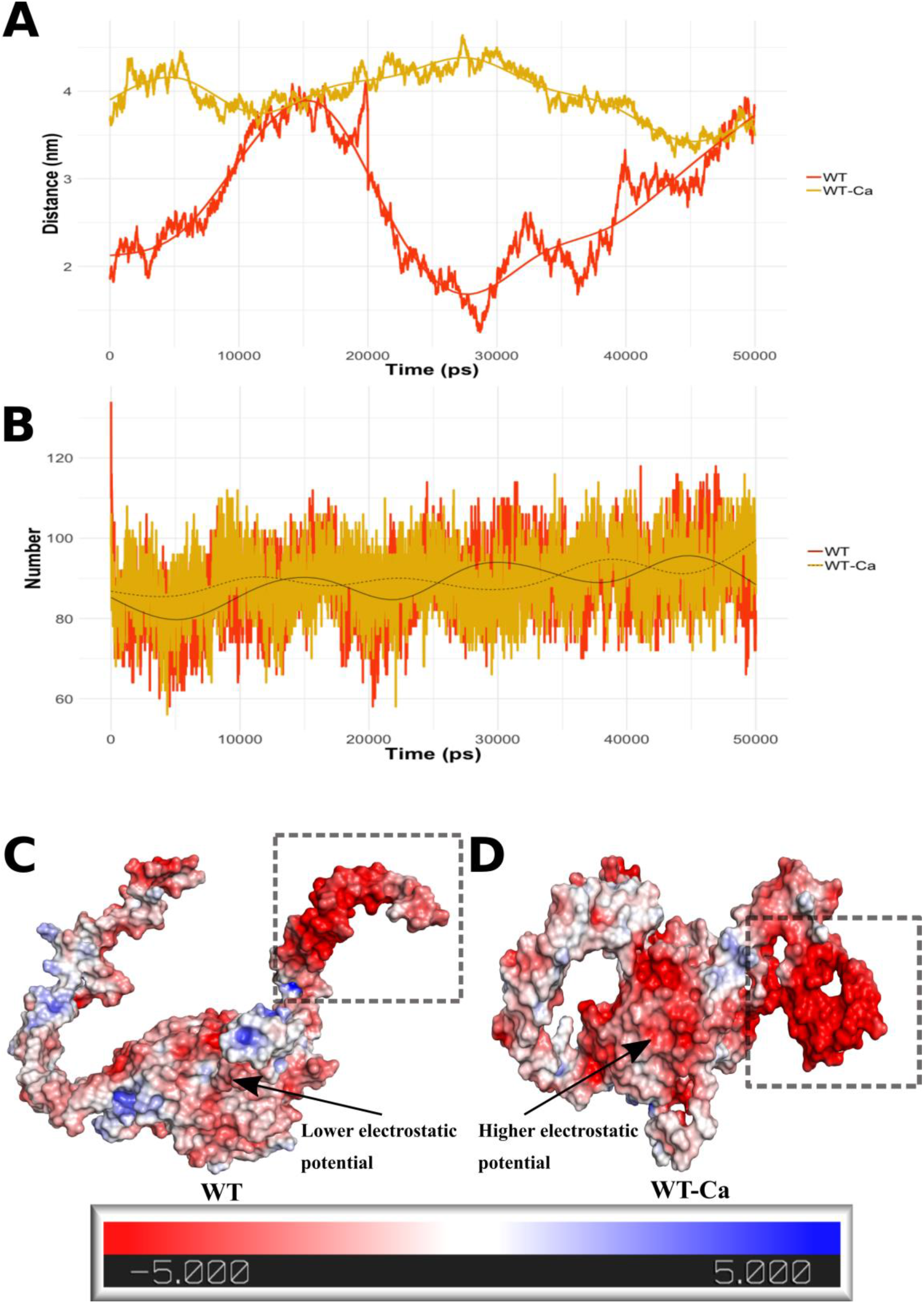
Comparative analysis of calcium dynamics and electrostatics between WT and WT-Ca. (A) minimum distance of ions from exon-9. Red color: distance of Na^+^ ions from WT. Golden: distance of calcium ions from WT-Ca. (B) number of intra-hydrogen bonds between WT and WT-Ca. Red: WT. Golden: WT-Ca. (C) electrostatic potential in WT. (D) electrostatic potential in WT-Ca. Red: higher electrostatic potential, blue: lower electrostatic potential. Areas shown with arrows and highlighted with dashed lines indicate differences in electrostatic charges.

Also, the mere presence of Na ions was not enough to effectively bind to the negative charges present in the C-domain compared to Ca ions while for mutated genes the electrostatic potential was too low for calcium ions to bind. Overall, the WT protein in the presence of calcium ions showed higher electrostatic interactions with calcium ions that proteins derived from mutated genes (Figure 6 and S6).

### Interaction of Exon-9 with Calcium Ions stabilizes the N-Domain and P-Domain

We aimed to examine whether the stabilization of the C-terminal domain could impart stability to the overall protein structure. To this end, we performed visual inspection of the 50^th^ ns structures from the simulated trajectory. The extended arm-like structure in the P-domain remains stable due to the close association with the N-domain. Residues Gly140, His241 and Ile242 from the N- and P-domains interacted with each other by forming hydrogen bonds after 30ns of simulation, thereby providing anchorage to the P-domain (Figure 8A,B). This interaction was possible because of the presence of the β-bend and β-bridge in WT-Ca while this was lacking in mutated counterparts.

**Figure 8:**
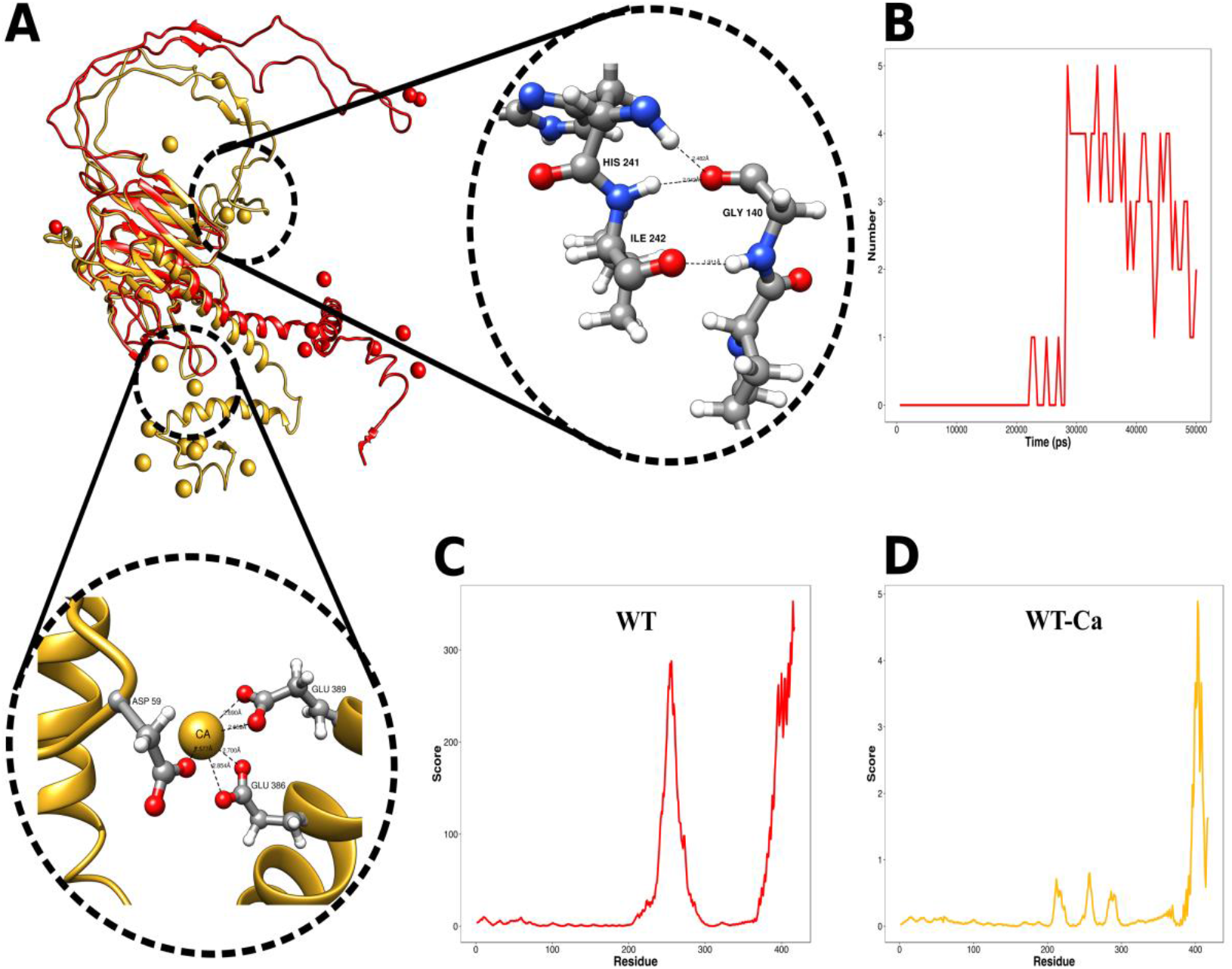
Interaction of various domains in presence of calcium ions. (A) key residues responsible for the interaction of N-domain to P and C domain. WT (red) has no domain interaction while WT-Ca (golden) undergoes structural transformation through the domain interaction. N-P domain is stabilized due to the interaction between Gly140with His241 and Ile242. N-C domain is stabilized due to the interaction of Asp59, Glu60, Glu386 and Glu389 in presence of Calcium ions represented in golden color. The red balls represent oxygen atoms while the white balls represent hydrogen atoms, the blue balls represent nitrogen and the grey color represents carbon backbone. (B) formation of hydrogen bonds between N-P domain throughout simulation. (C) and (D) represent the mobility of WT in presence and absence of calcium ions.

Additionally, the N-domain and C-terminal domains also interacted with each other in the presence of calcium ions. Asp59 and Glu60 of the N-domain interacted with Glu386 and Glu389 of the C-terminal domain suggesting that calcium ions not only stabilize the C-terminal region but also brings conformational changes to the C-terminal region in order to interact with the N-domain, thus providing overall stability. The 2D interaction plot between the above residues were cross-validated using Ligplot software [47] (Figure S7 A, B). The residue mobility scores for WT and WT-Ca indicated that the mobility of the residues of the P- and C-terminal domains in the WT was very high with a score of 300 while for WT-Ca the score was merely 5 suggesting a higher steadiness in WT-Ca (Figure 8C and D). Secondary structure comparison between WT and WT-Ca suggested that residues 138-140 and 241-242 hold a β-bend configuration in WT-Ca but not in WT. Moreover, the key residues Gly140-Ile242 undergo changes with time forming a β-bridge which makes the interaction more prominent. Furthermore, domain movement analysis by VMD in the form of porcupine plot described that the movement of the C-terminal domain in WT was inwards but outwards in WT-Ca as indicated by the arrows (Figure S8 A, B).

Next, we examined the exon-9 structural changes in WT and WT-Ca throughout the simulation period. Interactions of all the protein residues to calcium ions are shown in figure S9 and a full list of residues is shown in Table 3.

**Table 3:**
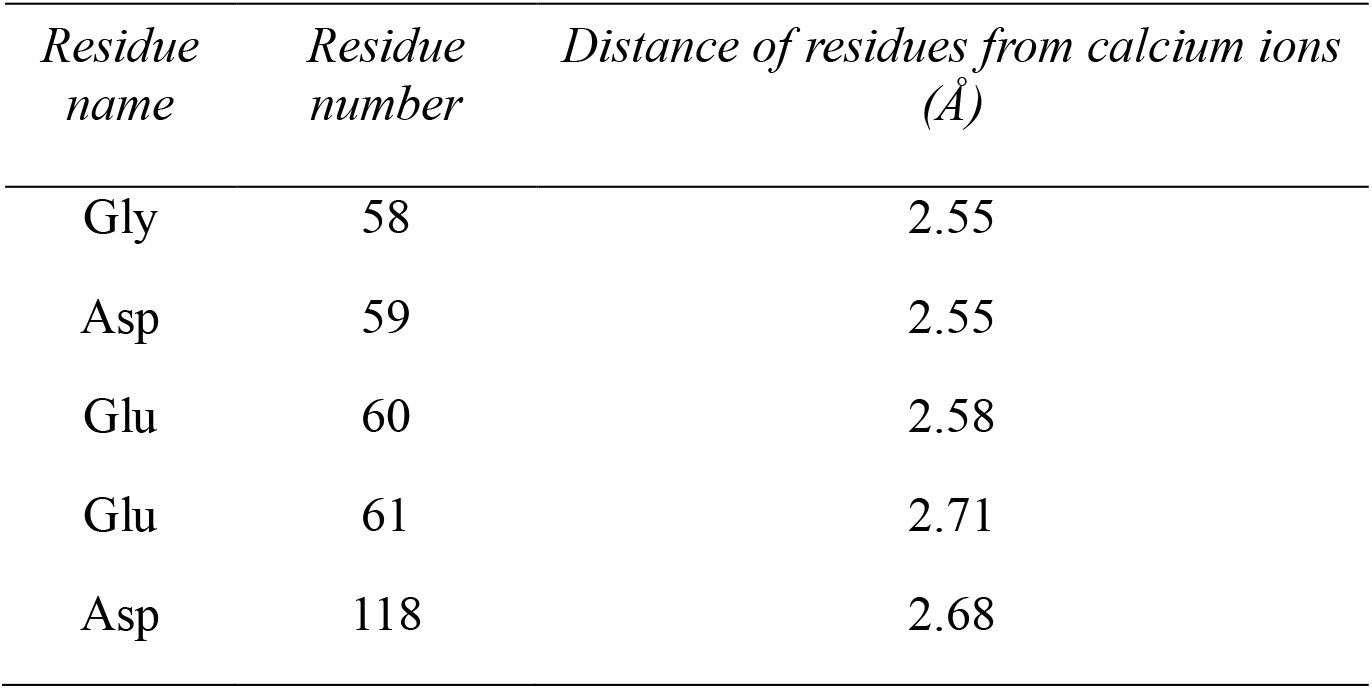
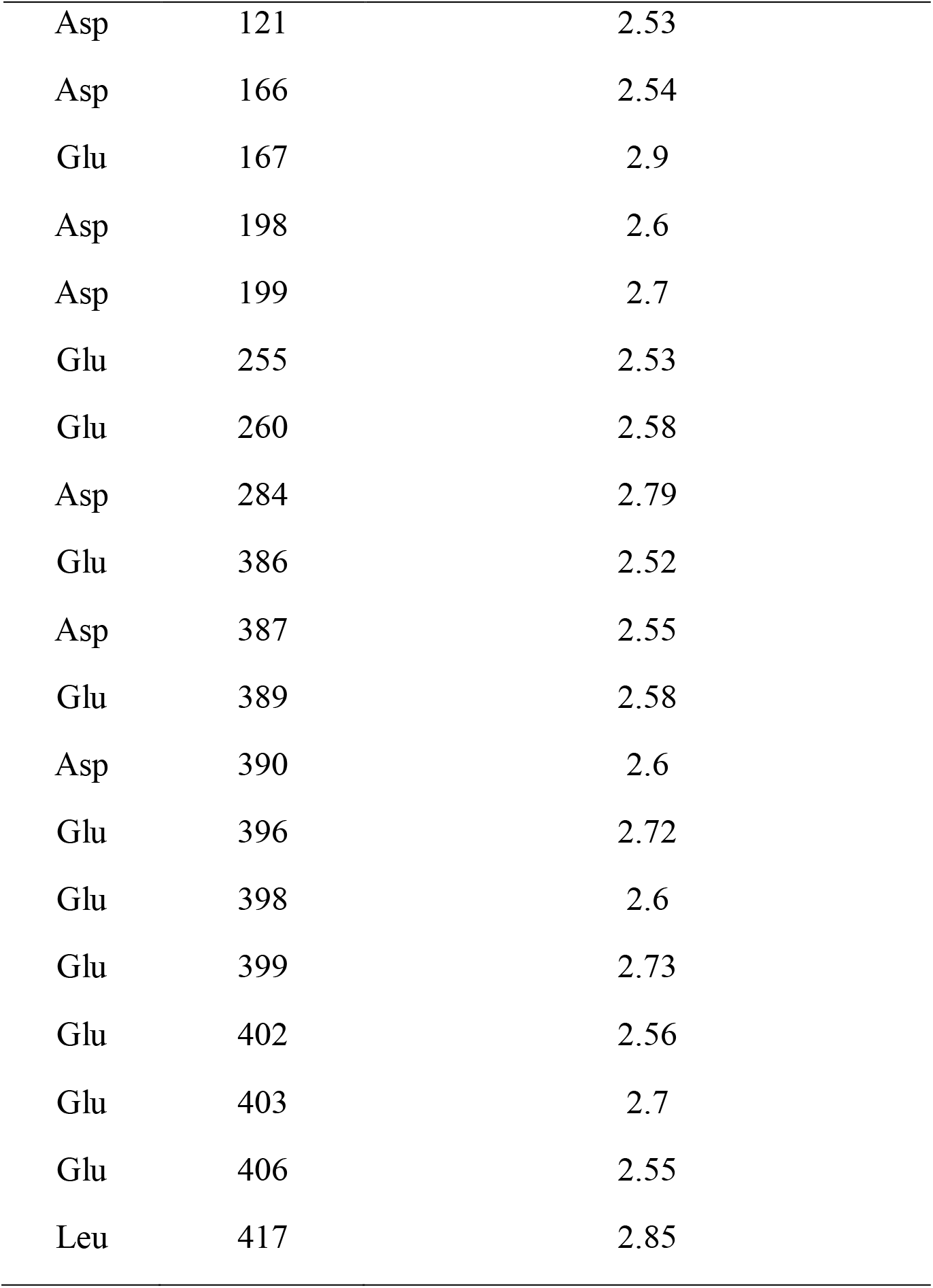
List of WT residues interacting with calcium ions

| *Residue name* | *Residue number* | *Distance of residues from calcium ions ($\text{\AA}$)* |
| --- | --- | --- |
| Gly | 58 | 2.55 |
| Asp | 59 | 2.55 |
| Glu | 60 | 2.58 |
| Glu | 61 | 2.71 |
| Asp | 118 | 2.68 |
| Asp | 121 | 2.53 |
| Asp | 166 | 2.54 |
| Glu | 167 | 2.9 |
| Asp | 198 | 2.6 |
| Asp | 199 | 2.7 |
| Glu | 255 | 2.53 |
| Glu | 260 | 2.58 |
| Asp | 284 | 2.79 |
| Glu | 386 | 2.52 |
| Asp | 387 | 2.55 |
| Glu | 389 | 2.58 |
| Asp | 390 | 2.6 |
| Glu | 396 | 2.72 |
| Glu | 398 | 2.6 |
| Glu | 399 | 2.73 |
| Glu | 402 | 2.56 |
| Glu | 403 | 2.7 |
| Glu | 406 | 2.55 |
| Leu | 417 | 2.85 |

Extracted static PDB files from the simulation trajectory at timeframes of 10ns, 30ns and 50ns expounded that exon-9 in WT was unstable while WT-Ca showed a very stable assembly (Figure 9). The majority of residues that interacted with calcium were either glutamic acid or aspartic acid. The two alpha-helices in exon-9 of WT-Ca maintained a stable conformation of ‘Λ-shape’ throughout the simulation. The two helices in exon-9 are separated by an angle of 70.57° while in WT-Ca by 54.44°. Moreover, in the absence of calcium, the two helices were separated by a distance of 38.73 Å while in the presence of calcium they were separated by 29.9 Å. The presence of calcium ions provoked the C-terminal region to be displaced towards the opposing sides as indicated by the angular displacement of the C-terminal region in WT and WT-Ca (Figure S10). This stability could only be attained due to the higher magnitude of interaction with calcium ions which holds the structure steadily.

**Figure 9:**
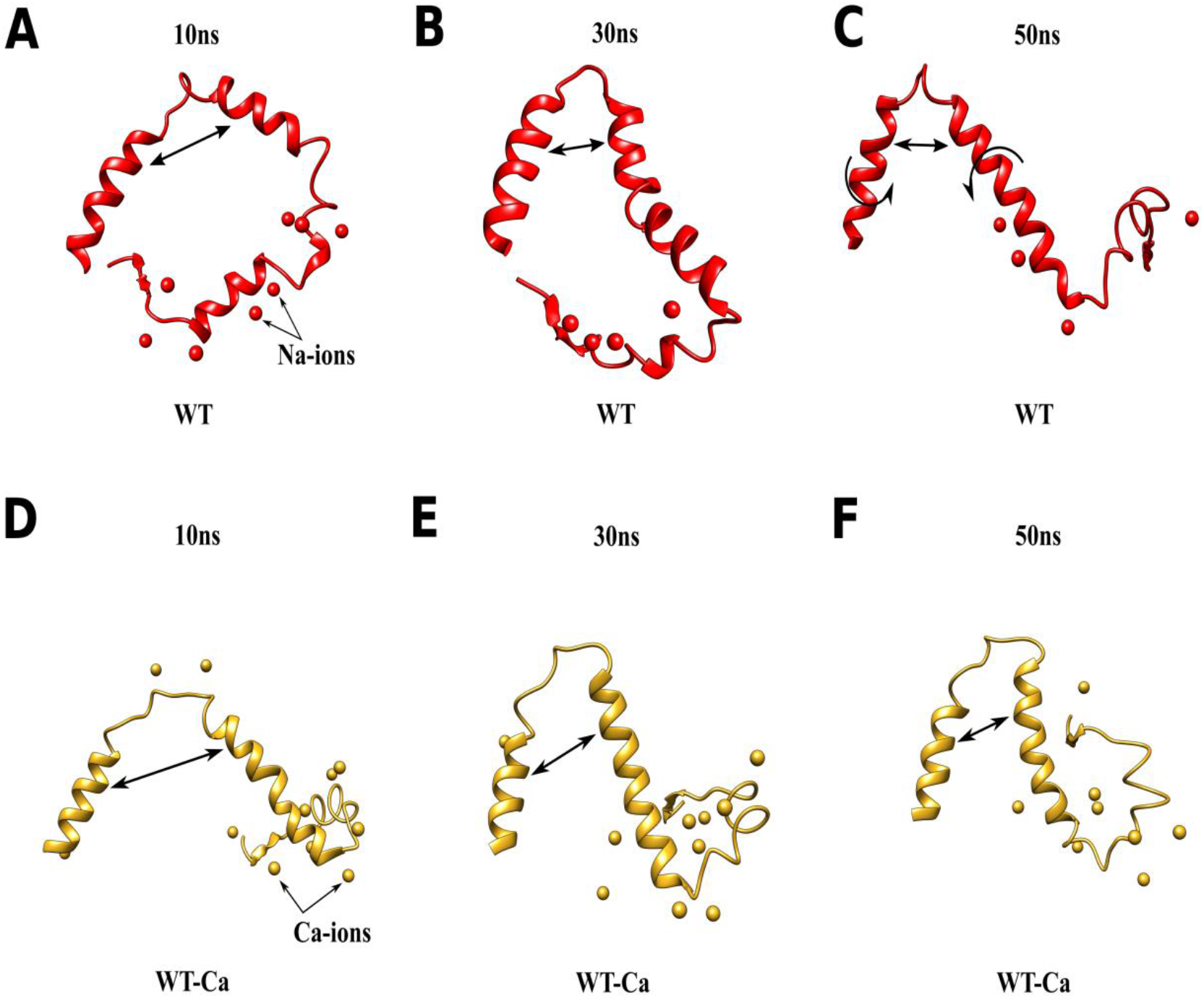
Time series Conformational changes in Exon-9 of WT and WT-Ca. (A-C) conformational changes and local transformations in exon-9 in the absence of calcium with time lapse of 10ns, 30ns and 50ns (D-F) conformational changes and local transformations in exon-9 in the presence of calcium with time lapse of 10ns, 30ns and 50ns.

### Exon-9 Mutation leads to Imbalance of Calcium Flux

We then sought to explore the effect of exon-9 mutations on protein structure *in vitro* in order to understand if there are any significant changes in their calcium regulation and fluxes. The transcriptional factor family Nuclear factor of activated T-cells (NFAT) is known for its involvement in calcium regulation [48, 49] and thus we first tested their mRNA levels. To our surprise, three members of the NFAT family members, namely NFAT 1, 3 and 5 were increased significantly in exon-9 mutated clones suggesting an imbalance of cellular calcium levels (Figure 10F). In order to validate the above mentioned hypothesis we explored the difference between WT and exon-9 KO cells using fluorescent microscopy. WT and exon-9 mutated cells were incubated for 30 mins with the Fluo4-AM probe and calcium signals were recorded using the 488 nm fluorescence channel. Both *CALR* exon-9 KO clones showed a sharp decrease of a spontaneous Fluo4-AM signal around the plasma membrane / extracellular region in comparison of WT. *CALR* has been found to localize in the extracellular region [50] and thus in the WT a higher calcium signal is observed as exon-9 can buffer and hold calcium ions while in the mutated exon-9 no such signal is detectable. To our surprise, there was a huge difference between the fluorescence intensities of calcium ions in the WT and mutated counterparts (Figure 10G). In sum, our *in-vitro* results showed that exon-9 plays a very crucial role in calcium binding and any mutation can drastically effect the cellular calcium levels.

**Figure 10:**
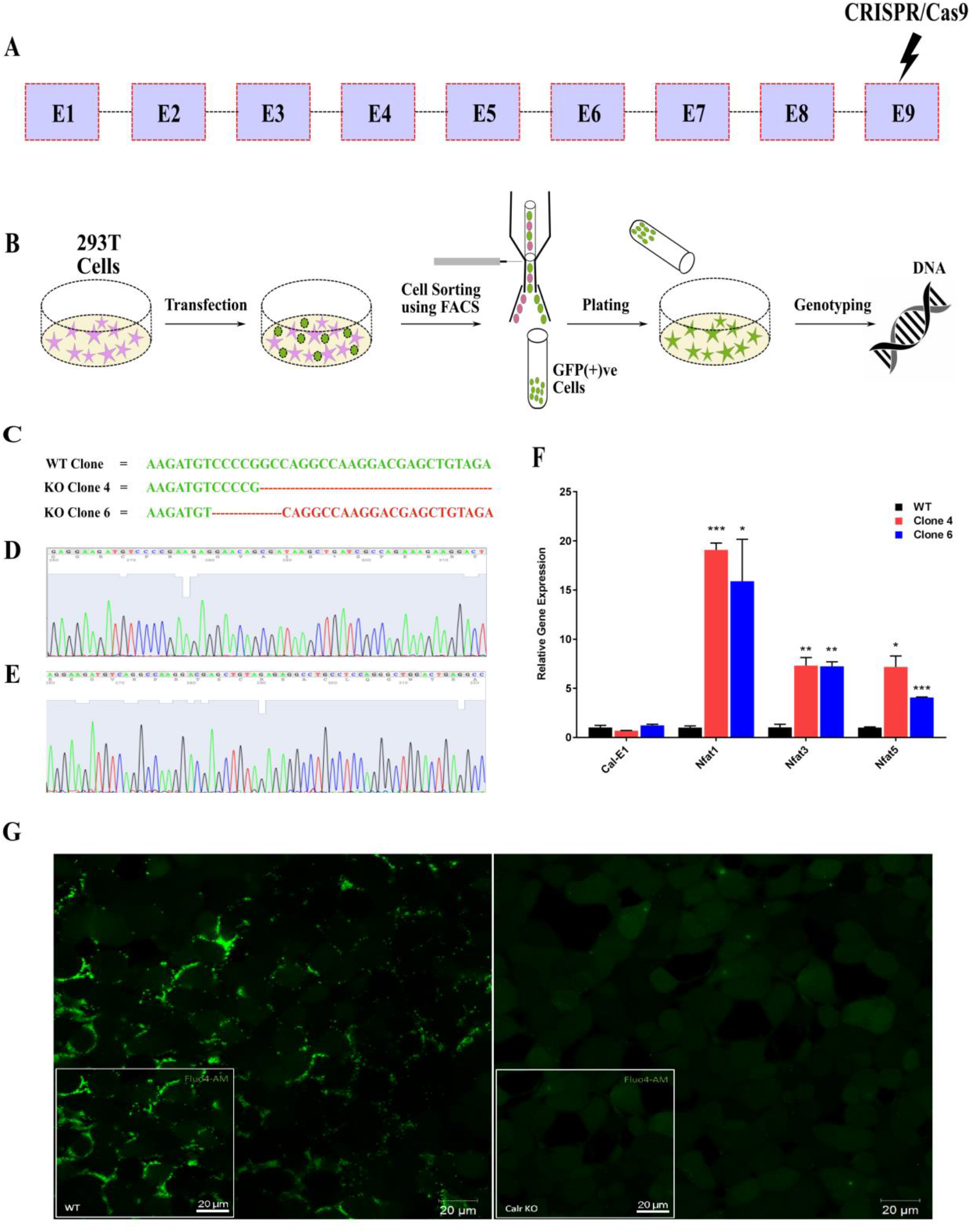
In-vitro results of WT and Mutated *CALR*. (A) Gene map of *CALR* (B) generation of *CALR* exon-9 mutations using CRISPR-Cas9 (C) sequence analysis of mutated clones taken under investigation (D)-(E) Sanger sequencing results of the mutated clones compared to WT (F) qPCR results of *CALR* Exon-1, *NFAT1, NFAT3* and *NFAT5* in WT and two mutated clones: clonec4 and clone 6 where WT is depicted in black color, clone4 in red and clone 6 in blue respectively (G) representative images of Fluo4-AM staining indicating the fluorescence intensity of calcium ions in WT and mutated counterparts.

## Discussion

Essential thrombosis (ET) belongs to a sub-category of myeloproliferative neoplasms (MPNs) which enhances the formation of blood clotting through an unknown mechanism. Studies have found that mutations in two genes are responsible for such disorder where Janus kinase 2 *(JAK2)* and CALR account for 50-60% and 40-50%, respectively. Much is known about the JAK2 mechanism while the role of CALR remains obscure although it was discovered as a susceptible gene a long time ago [51]. CALR is a highly conserved protein bestowed mostly with negative charged amino acids such as aspartic acid (Asp) (13.2%) and galutamic acid (Glu) (12.9%) while exon-9 accounts for 21.2% and 37.90% (Figure S5D) of the two amino acids, respectively, a two-fold increase in Asp and three-fold in Glu. MPN patients encompass heterogeneity specifically in exon-9 leading to loss-of-function mutation and thereby generating a novel C-terminal sequence. The wild type CALR protein localizes in the ER to ensure proper folding of glycoproteins as well as to maintain calcium homeostasis [52]. Unavailability of a three-dimensional structure provoked us to model the full-length structure in order to visualize the overall structural dynamics in the absence and presence of calcium ions. As discussed by Abreusan et al. [53], an alpha helix provides more stability and robustness to proteins endorsing the fact that three out of four alpha-helices of CALR are encoded by in exon-9 albeit the mutation frequency is highest in exon-9. Moreover, the protein derived from exon-9 exhibits a helix-loop-helix structure and almost all of the INDEL mutations in humans start from Gln365 [5] which lies in the loop region while variations among different species start from E383 which lies in the helix region (Figure S1). The INDELs lead to the distortion of the C-terminal domain into the P-domain making a closed conformation due to the loss of structural integrity in both mutated isoforms (Figure 2B-C & 4B-C). These mutated structures are more stable due to less fluctuations in their C terminal region, also evidenced by their kinetic energy (KE) and they could readily interact with other proteins due to higher potential energy (PE). However, in the presence of calcium ions the WT protein attains conformational stability and could interact more readily with other proteins indicating the need of calcium ions for proper functioning as indicated by higher KE and PE (Figures 2 D-E, 3, 4D-E and Table S4). This is in line with the previous studies performed by Akari et al. [54] as the CALR mutants could interact more firmly with the thrombopoietin (TPO) receptor activating JAK2 and its downstream molecules such as mitogen-activated protein kinase 3 (MAPK3 or ERK1) and Signal transducer and activator of transcription 5 (STAT5). The binding of mutant CALR to the TPO receptor could presumably activate platelet production; which is also a common phenotype in ET patients. A similar study was performed by Pronier et al. [55] as they traced the interacting partners of mutated isoforms identifying friend leukemia integration 1 (FLI1) - a transcriptional regulator, as the key candidate that activates the TPO receptor and increases megakaryocytes formation which could be responsible for higher platelet production. Interestingly, Elf et al. had shown that mutant CALR binds to the TPO receptor which triggers MPN progression [56]. Thus, the modified C-terminal region in CALR proteins derived from mutant gnes might play a vital role in such interaction. Moreover, a direct study by Kollmann et al. [57] states that mutant CALR facilitates megakaryocyte production by MAPK signaling requiring calcium ions [58, 59]. Apart from this, a recent study from Shide et al. [60] reported that homozygous mutations of *CALR* in mice result in mild thrombocytosis suggesting that a mutant C-terminus can undergo numerous modifications depending on the position of the mutation and involved amino acids. A possible explanation could be that the *CALR* sequence in the C-terminus varies between human and mouse, giving rise to a different mutated C-terminal region with different binding propensities. Comparing human and mouse amino acid sequences, a few important amino acid substitutions like E60L in the P-domain, E402del in the C-terminal domain and V409S could play a vital role as in humans these residues are critical for providing stability in presence of calcium (Figures 1, 8, S7B and Table3). These variations in amino acids could differ in calcium binding strength which could give rise to a non-interacting and non-functional C-domain.

The presence of calcium in the system provided clues that only the WT can bind to calcium ions due to the electrostatic interactions while INDELs could not as they lacked the negative C-terminus. The negatively charged environment in exon-9 is able to compensate for the higher calcium load and maintained its conformational stability (Figures 5, 8). 27.5% of total calcium ions remained bound to the C-terminal region while only 6.7% (Figure 8) accounted for the WT indicating that the C-terminal region is preferred by Ca^2+^ ions and not Na^+^ ions. Our study also shed light on demonstrating that a monovalent ion like Na^+^ is not enough to counter-balance the negative potential of the C-terminal region. Further studies are required in order to understand whether other divalent or trivalent ions could also provide such stability and interaction. An important observation in the current study is that conformational changes were not limited to exon-9, but the stability in exon-9 modulated the distally located P-domain in response to calcium binding. The P-N domain attained its structural stability due to the hydrogen bonds between His241-Gly140 and Ile242-Gly140 while the interaction of the C-N domain was contributed by the acidic residues Glu386, Glu389, Glu60 and Asp59 (Figure.8).

One of the key structural changes observed between WT and mutated proteins is the displacement of the second helix in the C-terminal domain (Figure S10) where helix 1 and helix 2 of the WT makes an angular displacement of 70.57° and are separated by a distance of 38.73Å while in WT-Ca the two helices are much closer to each other with a distance of 29.99Å and an angular displacement of 54.44° (Figure S10). In the absence of calcium ions, WT and mutated counterparts reflected an opposite pattern of folding. WT remained in open configuration while mutated ones in closed indicating that the proteins derived from mutants have a very different architecture of amino acids which could trigger a cascade of reactions involved in calcium signaling as well as platelet formation causing diseases like ET. Although the mutations increased the structural stability, they lose their functionality as loss of the KDEL motif from the exon-9 would remove the ER signature and thus they could no longer function as an ER chaperone and would localize to other cellular compartments. This loss of the ER retention signal would increase the calcium potential thereby disrupting the natural pH of 7.2 in the ER [61] and increasing an acidic ER stress. Moreover, recently it was reported that CALR could buffer ~50% of ER calcium suggesting a very high level of calcium influx could be generated due to improper buffering [9].

CALR acts as a double edged sword. In one way, WT CALR plays a crucial role in buffering cellular calcium load due to the presence of a highly acidic C-terminal domain, and in return calcium ions provide structural stability to CALR by balancing the electrostatic charges. In contrast, a mutated CALR structure has a higher binding affinity for the TPO receptor that activates platelet-related transcription factors, as calcium plays a vital role in the production of platelets. In a wider sense, the current study touches upon the possible role of calcium ions in the formation of ET, although the sequences in the middle of exon-9 are evolutionary not conserved (Figure S1). One possible explanation could be that exon-9 was merged with CALR by natural selection to maintain calcium load and homeostasis only when calcium ions were available for uptake and utilization. The higher negative charge in exon-9 compared to other exons would help to effectively utilize calcium ions by neutralizing and balancing calcium charges. Exon-9 structural dynamics shows that gaining and loosing amino acids in the derived proteins by mutations significantly affects the structure which in turn would affect function (Figure S11). Overall, the findings from this study suggest a loss-of-function mutation in exon-9 of *CALR* which drives the onset of essential thrombocythemia through the imbalanced calcium load. Our result provides new structural insights into the working mechanism of intrinsically disordered *CALR* exon-9. We identified key residues involved in domain stability. Prospects lie in conducting experiments for pH-dependent simulations in order to examine the correct physiological pH for CALR to be functional in the presence of calcium. It is also worth investigating possible key residues involved in the interaction between mutant CALR and the TPO receptor which will help in the screening of inhibitors to act as an antagonist for JAK2 activation.

Calcium also acts in cell-to-cell communication and any change in the calcium level would trigger a response [62] in controlling a spectrum of cellular processes through calcium signal transduction [63]. Calcium ions are also essential for protein function as they can change protein shape and electrostatic charge distribution [18]. Hence, the protein derived from exon-9 being highly electronegative in nature, can easily bind and stabilize the charge of positive calcium ions. Exon-9 mutations imbalance the calcium homeostasis as evidenced from results highlighted in Figure 10 (G). WT cells have much higher calcium levels as CALR binds and sequesters calcium ions, while in contrast, mutant CALR is not able to hold calcium ions leading to very low cellular calcium levels. This is in accordance with a previous study where the authors showed that non-endoplasmic reticulum based CALR can affect calcium signaling [9]. Decreased extracellular calcium level hampers cell communication and finally leads to slower cell growth as imbalance of calcium ions could induce the G0 phase of the cell cycle [64]. This is in line with our data on slower cell proliferation of exon-9 KO 293T cells (data not shown). We also observed an increased mRNA level of *NFAT1,3* and *5* in exon-9 KO 293T cells, probably because cells are in a hypo-calcium state and thus requiring a higher calcium flux to maintain cellular homeostasis (Figure 10F), which remains to be further investigated.

## Conclusion

Although 40-50% of ET is caused due to the mutations in the *CALR* gene, the exact mechanism remains vague. To understand the mechanistic insights of CALR dynamics in disease progression and etiology, this study employed full length structures of three isoforms of *CALR* and investigated the dynamics and transitions based on standard MDS. Secondly, we also studied the effect of calcium dynamics on the isoforms of the proteins to understand the calcium dynamics in regulating CALR. RMSD, Rg and RMSF between the isoforms of CALR revealed that the WT protein attains a highly unstable conformation compared to the two mutational isoforms. Two domains, namely the proline-rich P-domain and C-terminal domain displayed disordered movements. However, in the presence of calcium ions, the WT protein surmises a more compact and stable conformation than its mutated counterparts as a result of lower residue fluctuations in P and C-terminal residues. Electrostatic effects mediated by the highly negative C-terminal domain attract free calcium which significantly decreases the enhanced fluctuations present in the C-terminal region. The affinity of the C-terminal region towards calcium ions further results in providing enhanced stability to the N-terminal region as well. Thus, the presence and absence of calcium ions display contrasting behavior in the WT and mutated CALR. The mutated isoforms lack the electrostatic ability due to the absence of negative amino acids resulting in ineffective binding to calcium ions thereby perturbing the cellular calcium flux. We uncovered that the P-domain and C-terminal domain undergoe structural changes in the presence of calcium ions, thereby maintaining structural stability which can then interact with other proteins. The negative potential of the exon-9-drived protein alone is crucial for influencing stability and calcium homeostasis. Finally, Fluo4-AM staining indicated a huge difference in calcium intensity between WT and exon-9 KO cells. Our results should facilitate an understanding of ET formation via CALR mediated calcium signaling and might assist in finding therapeutic interventions for treating ET.

## Materials and Methods

### Sequence analysis

The protein sequence of *CALR* with the accession number P27797 was retrieved from Uniprot database (UniProt, 2015) in Fasta format, and the coding DNA sequence (CDS) with ID of ENSP00000320866.4 was retrieved from the Ensembl database [24]. According to a previous annotation [5], a 52 base pair deletion (Del) and 5 base pair insertion (Ins) were manually manipulated in the coding region of exon-9 in-frame to mimic *CALR* mutations. The secondary structures of the wild type and mutated *CALR* were examined by the PDBsum web server [25]. CALR domains were characterized by using the Pfam [26], InterProScan [27] and Uniprot web-servers, and the physiochemical property of *CALR* was examined with the Protparam tool [28].

### Structural characterization

Currently only a limited part of the *CALR* structure could be resolved [29]. The I-TASSER package [30] was used to construct the full-length CALR structure through homology modelling based on the multiple threading alignment by LOMETS [31] and the iterative TASSER [32] assembly simulations. The *CALR* mutant structures were constructed similarly based on the INS and DEL sequences. The built models were evaluated using the PROCHECK server [33], VERIFY3D [34], and ERRAT web servers [35]. The best-performed structures were firstly energy minimized using the steepest descent method to remove bad contacts within atoms using the Swiss-PDB viewer [36] and then subjected to a molecular dynamics (MD) simulation.

### Molecular Dynamics Simulation

To characterize *CALR* and identify the role of Ca^2+^, molecular dynamics (MD) simulations on the WT (Wild Type) form as well as two mutant-forms namely Ins (inserted mutant) and Del (deleted mutant) together with the protein-Ca complexes WT-Ca, Ins-Ca and Del-Ca were performed using the GROMACS 4.6.5 package [37] combined with Amber99sb-ildn force field [38]. Each model was immersed into the center of a triclinic box of extended simple point charge (SPC-E) water molecules [39] with a margin distance of 10.0 Å. Periodic boundary conditions were applied to all directions. To keep the system in charge neutrality, Na^+^ and Ca^2+^ counter ions were added to the WT, Ins, and Del forms making two experimental groups – sodium bound group (WT-Na, Ins-Na, and Del-Na) systems and calcium bound groups (WT-Ca, Ins-Ca, and Del-Ca) systems, respectively. 30000 steps of steepest descent energy minimization (EM) and 20000 steps of conjugated gradient EM were conducted in order on the constructed models. Then each system was gradually heated from 0 K to 300 K under conserved number of moles, volume and temperature (NVT) ensemble over a period of 500 pico-seconds (ps), during which a weak constraint of 10 kcal·mol^-1^·Å^-2^ was imposed onto the protein. Afterwards, each model was subjected to an equilibrium simulation for 1000 ps by removing all constraints. Finally, a production simulation for each model was conducted under conserved number of moles, pressure and temperature (NPT) ensemble. The reference temperatures were kept at 300K, the LINCS algorithm (Hess et al., 1997) for bond constraints was used. The MD trajectory was recorded every 2 ps. VMD [41] and Chimera [42] were applied for visualizing and analyzing the MD results.

### Cell Culture

HEK-293T cells were cultured on cell culture dishes (Corning) and maintained in cell culture medium comprised of Dulbecco’s modified Eagle’s medium (Hyclone) supplemented with 10% fetal bovine serum (FBS) (Sigma-Aldrich) and penicillin-streptomycin antibiotic solution (Sigma-Aldrich) under humidified conditions at 37 °C, 5% CO_2_.

### CRISPR plasmids preparation

The CRISPR-Cas9 plasmid pSpCas9(BB)-2A-GFP (PX458) (Addgene plasmid #48138) was used for generating the knockout mutants. Custom designed guide RNAs were synthesized (Generay, China), annealed and ligated into BbsI digested sites. The ligated products were transformed into chemically competent DH5α *E. coli* cells (Transtaq, China). The ligated cells were then plated in Ampicillin selection plates (100 μg/mL ampicillin) and incubated at 37 °C for 12–14 hours. Single and isolated colonies were picked and expanded in LB medium containing ampicillin. Plasmid DNA was isolated using the Taraka Mini Plasmid Extraction Kit and insertion of guide RNAs was verified by Sanger sequencing using the universal forward primer for the human U6 promoter (5’ GACTATCATATGCTTACCGT 3’) (Biosune, China).

### Generation of a CALR mutant in HEK-293T cells

To generate a *CALR* exon-9 mutant cell line, HEK-293T cells were first cultured in 24 well cell culture plates (NEST, China). The protocol by Ran at al. was used to generate the exon 9 KO cells [43]. After the cells attained 70% confluency, 500 ng CRISPR-Cas9 (PX458) plasmid having the right sgRNA insert (Table S1) was transfected along with transfection reagent Lipofectamine 2000 (Sigma) at a 1:3 DNA to reagent ratio. Transfection medium was changed after 12 hours from transfection to the complete medium. Cells were harvested for fluorescence-activated cell sorting (FACS) (BD Biosciences) after 24 hours of incubation. As the plasmid was fused with GFP, GFP(+ve) cells were sorted in 96 well plate (Corning) and a total of 200 cells were sorted for each guide RNA. The medium was changed every five days and single clones were expanded for 21 days. The clones were picked according to confluency for further expansion, DNA was extracted using the Takara Universal Genomic Extraction Kit (TakaraBio) and was sent for sequence verification (Biosune, China).

### RNA Extraction and qPCR

Total RNA was extracted using the TRIzol reagent (Invitrogen) according to the manufacturer’s instructions. The RNA concentration was measured using a NanoDrop 8000 spectrophotometer (Thermo Scientific). 1 ug RNA was reverse transcribed using a PrimeScript RT-PCR kit (Takara Bio) by following the manufacturer’s instructions. The cDNA was then employed for quantitative polymerase reaction (qPCR) using a SsoAdvanced™ SYBR^®^ Green Supermix (Bio-Rad, USA) with denaturation at 95 °C for 30 s, followed by 40 cycles of denaturation at 95 °C for 15 s, and annealing at 60 °C for 30 s in a CFX96 System (Bio-Rad). The primer sequences are listed in supplementary Table S1.

### Calcium imaging

Cells were seeded on a glass cover slip chamber with 200 μl complete medium. The cells were grown until 70% confluency and incubated with the calcium indicator Fluo-4 AM (1μM) (Invitrogen, US) for 30 minutes at room temperature. Images were captured using an inverted Zeiss LSM710 confocal microscope with a 40x oil immersion objective. A wavelength of 488 nm was used to excite the samples and fluorescence emission was collected at > 505 nm for single color calcium imaging.

### Statistical Analysis

Statistical analysis between two groups was performed by Student’s T-test using GraphPad Prism version 7 for Windows (GraphPad Software, La Jolla California, USA). A p-value of lower than 0.05 was considered significant.

## Abbreviations

ABPS: Adaptive Poisson-Boltzmann Solver
CALR: Calreticulin
Del: Deleted
ET: Essential Thrombocythemia
ER: Endoplasmic Reticulum
INDEL: Insertion and Deletion
Ins: Inserted
KE: Kinetic Energy
KO: Knock-out
MPN: Myeloid Proliferative Neoplasm
MDS: Molecular Dynamics Simulation
PE: Potential Energy
RMSD: Root Mean Square Displacement
RMSF: Root Mean Square Fluctuations
Rg: Radius of Gyration
WT: Wild type

## Data availability

Source data are provided for all experiments. Other data that support the findings of this study are available from the corresponding author, upon reasonable request.

## Acknowledgments

The authors would like to thank Institute of Aging Research, Hangzhou Normal University for providing research facilities. We thank Dr. Gabriele Saretzki of Newcastle University for comments on the manuscript. Amit Jaiswal would also like to thank Ram Manohar Jaiswal and Pratima Jaiswal for their constant support throughout this research.

## Author Contributions

Conceptualization: Amit Jaiswal

Formal analysis: Amit Jaiswal, Zhiguo Wang

Figure Preparation: Amit Jaiswal

Funding acquisition: Zhenyu Ju, Zhiguo Wang

Investigation: Amit Jaiswal, Zhiguo Wang, Xudong Zhu, Zhenyu Ju

Methodology: Amit Jaiswal, Zhiguo Wang

Project administration: Amit Jaiswal, Zhiguo Wang, Zhenyu Ju

Writing original draft: Amit Jaiswal

Writing review & editing: Zhiguo Wang, Xudong Zhu, Zhenyu Ju

